# Meeting Measurement Precision Requirements for Effective Engineering of Genetic Regulatory Networks

**DOI:** 10.1101/2021.10.10.460840

**Authors:** Jacob Beal, Brian Teague, John T. Sexton, Sebastian Castillo-Hair, Nicholas A. DeLateur, Meher Samineni, Jeffery J. Tabor, Ron Weiss, the Calibrated Flow Cytometry Study Consortium

## Abstract

Reliable, predictable engineering of cellular behavior is one of the key goals of synthetic biology. As the field matures, biological engineers will become increasingly reliant on computer models that allow for the rapid exploration of design space prior to the more costly construction and characterization of candidate designs. The efficacy of such models, however, depends on the accuracy of their predictions, the precision of the measurements used to parameterize the models, and the tolerance of biological devices for imperfections in modeling and measurement. To better understand this relationship, we have derived an *Engineering Error Inequality* that provides a quantitative mathematical bound on the relationship between predictability of results, model accuracy, measurement precision, and device characteristics. We apply this relation to estimate measurement precision requirements for engineering genetic regulatory networks given current model and device characteristics, recommending a target standard deviation of 1.5-fold. We then compare these requirements with the results of an interlaboratory study to validate that these requirements can be met via flow cytometry with matched instrument channels and an independent calibrant. Based on these results, we recommend a set of best practices for quality control of flow cytometry data and discuss how these might be extended to other measurement modalities and applied to support further development of genetic regulatory network engineering.

## Introduction

In mature fields of engineering, most design work is carried out “virtually” using quantitative numerical models to efficiently explore design space. Physical prototypes, on the other hand, are primarily used to validate models and designs and to refine predictions, rather than as a part of the search for some viable design. Once the goals of an engineering project reach even a moderate degree of complexity, the modeling and simulation of a proposed design is much faster and cheaper than the construction of a physical prototype, but such models are only useful if they are sufficiently accurate to reliably predict whether the physical instantiation is likely to satisfy the engineering goals. For a synthetic biology project, the number of potentially viable designs rapidly exceeds the number of combinations that can be readily brute-forced with high-throughput screening methods^1^. Moreover, high-throughput screening methods are still effectively out of reach of all but the most well-financed labs, meaning that most synthetic biology practitioners are only able to build and test a small number of constructs in each iteration of their engineering cycle.

Quantitative predictive modeling should allow practitioners to more efficiently explore design space, increase the likelihood that a physical prototype meets design goals, and thus decrease the barriers to entry for synthetic biology engineering. The efficacy of such predictions involves three components: accurate biological models, precise measurements providing parameters for the models, and biological devices with some degree of tolerance for imperfection in modeling and measurements. Recent years have seen much improvement in all three of these areas, including models that provide good accuracy in predicting certain compositional behaviors of engineered genetic constructs (e.g.,^2–10^), protocols for reproducibly precise measurement of genetic construct behavior^11–13^, and families of biological regulatory devices with improved performance (e.g.,^6,8,10,14–20^). Despite all of these advances, however, effective predictive engineering with genetic constructs remains extremely difficult and is only currently done under specific narrow circumstances.

This difficulty likely stems at least partly from persistent shortcomings in measurement reproducibility and precision. But just how reproducible do measurements actually need to be? Moreover, can we achieve a sufficient level of precision with available methods, and what best practices can help to ensure data actually attains the desired precision? To shed light on these questions, we derive a mathematical bound, which we call the *Engineering Error Inequality*, that quantifies the relationship between predictability of results, model accuracy, measurement precision, and device characteristics. We then apply this bound to estimate requirements for precision for measurements used to engineer genetic regulatory networks. Finally, we use data from an interlaboratory study to validate that these requirements can be met via flow cytometry with the aid of matched instrument channels and an independent calibrant. Based on these results, we recommend a set of best practices for quality control of flow cytometry data and discuss how these results might support more effective engineering of genetic networks.

**Table 1:** Definitions of key mathematical symbols used in this paper.

| Name | Definition |
| --- | --- |
| $v$ | a property value for a design |
| $r$ | the range $[r_-, r_+]$ of specification-satisfying values for $v$ |
| $\hat{v}$ | estimated value of $v$ |
| $\hat{r}$ | predicted range of $r$ |
| $\mu$ | geometric standard deviation of measurement error (fold) |
| $m$ | geometric standard deviation of model error (fold) |
| $p_{err}$ | probability a design instantiation fails to satisfy its specification |
| $p_s$ | probability a design instantiation succeeds in satisfying its specification |

## Results

### The Engineering Error Inequality

We propose that a conservative approximation of the difficulty of engineering a genetic construct can be made by considering a simple “atomic” act of engineering decision-making: checking whether a property value υ of a proposed system falls into a range of viable values *r* = [*r*_−_, *r*_+_]. Practically every choice in genetic construct design can be interpreted in terms of one or more such evaluations, for example:

- Will a given promoter drive production of an enzyme to a high enough level to support the intended reaction to generate a desired product? Here, υ is promoter activity and *r* is the range of activity levels that will result in the intended production rate.
- Is the high signal supplied to a genetic inverter sufficient to suppress its output below a desired level of expression? Here, υ is the input signal level and *r* is the range of input levels that will produce outputs below the threshold.
- Will a given chemical sensor detectably vary its output when exposed to the chemical at various concentration levels of interest? Here, υ is the minimum difference in outputs produced by the target concentrations and *r* is the range of output differences that can be detected.

The full details of any such engineering process will typically be much more complex. Most systems will involve more than one decision and it may be difficult to actually achieve a value in the viable range. The basic ability to check whether a value falls in a desired range, however, is a necessary precondition for any such more sophisticated decision-making. This implies an inequality relation (per details in the derivation given in the Methods), such that any metric quantifying the difficulty of evaluating a single value check is also a lower bound on the difficulty of evaluating any other design decision.

If we know the actual values of υ and *r*, then making such a check is trivial. In practice, however, uncertainty enters the decision-making process because we cannot directly access the true values of either υ or *r*. Instead, we must work with estimated values. Experimental characterization produces an estimated value 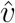 that varies from the unknown true value υ. Applying predictive models produces an estimated range 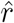 that varies from the unknown true value *r*. In the context of the engineering of genetic regulatory networks, gene expression distributions are typically log-normal ^21^ (or a close approximation^22^ thereto). As a consequence, the errors of measurements and model predictions are typically well-described in terms of ratios, rather than differences. In this case, we can thus describe the error of measuring υ (i.e., for which precision gives a lower bound) as the geometric standard deviation *μ* of values of 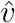. Likewise, model accuracy may be described as the geometric standard deviation *m* of predictions of the magnitude or bounds of 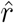. Figure 1(a) provides a diagram illustrating this relationship.

**Figure 1:**
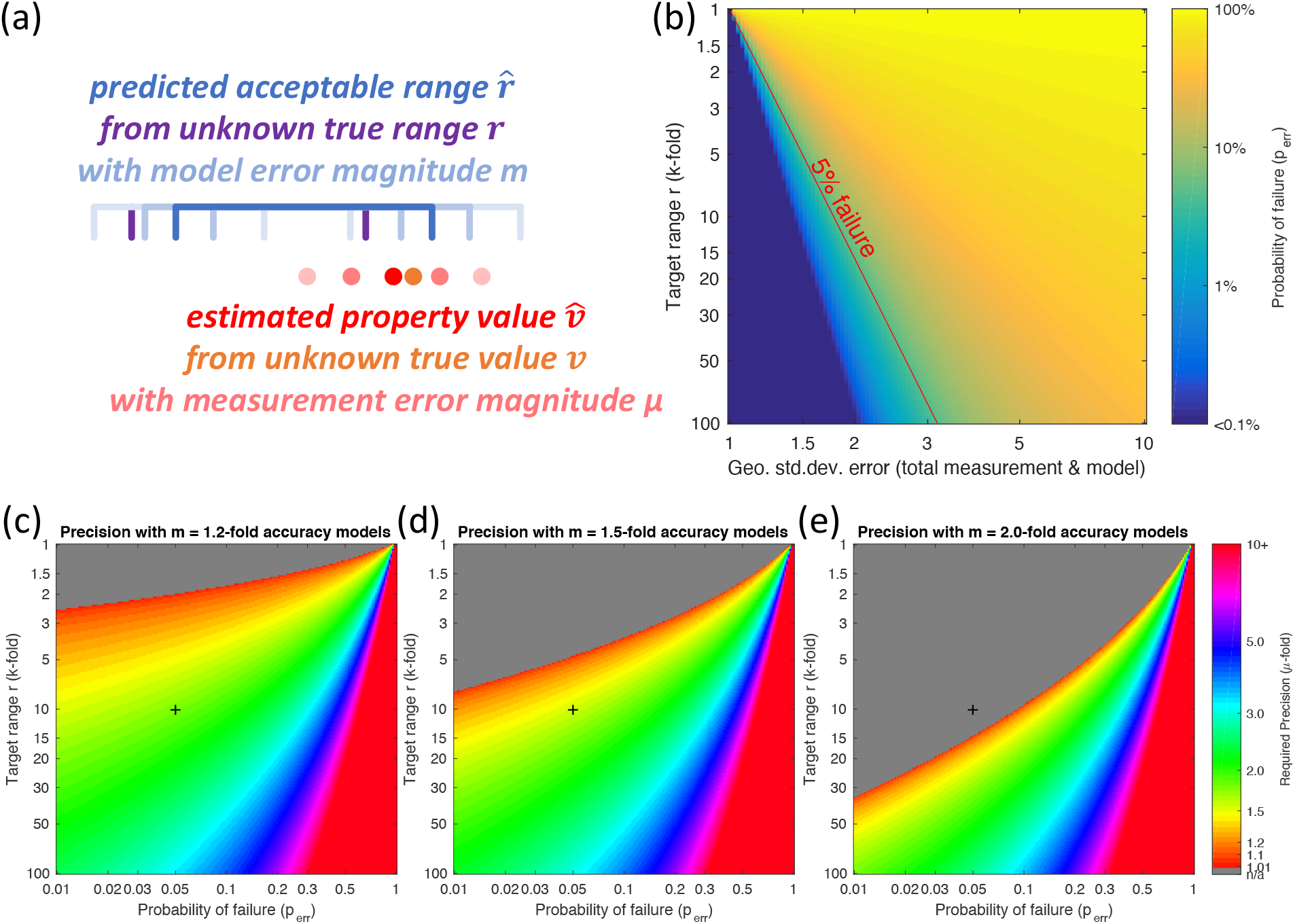
(a) Minimum requirements for effective engineering can be estimated by considering the effect of errors on an “atomic” engineering decision: checking whether an estimated property value 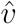 (red) lies within the estimated range 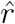 (blue) that can correctly realize an engineering specification, given a measurement error distribution of magnitude *μ* (light red) estimating the property value and a model error distribution of magnitude *m* (light blue) in predicting the acceptable range. The relationship of these values determines the probability that the true value υ (orange) actually falls within the true range *r* (purple); in the illustrated case, they do not, and the design will fail. (b) Minimum probability of failure with respect to target range and total error (combining both measurement error and model error); red line highlights the typical heuristic significance cutoff *p*_*err*_ = 0.05. (c, d, e) Examples of the measurement precision required to hit a target range with no more than given a probability of failure, given a model geometric standard deviation of 1.2-fold (c), 1.5-fold (d) or 2-fold (e). Black cross highlights the example of a 10-fold range and 5% chance of failure, which requires approximately 1.7-fold measurement precision with 1.2-fold model error, 1.5-fold precision with 1.5-fold model error, and is not possible with 2-fold model error.

For example, consider the case given above of selecting a constitutive promoter to drive production of an enzyme to an expression level υ within a range *r* of expression levels that will support the intended reaction. Here, the measured expression level 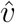 might be an average protein expression level of 3.5*e*4 molecules/cell to a precision *μ* of 2.5-fold standard deviation (i.e., 95% chance of falling within a range of 5.6*e*3 to 2.2*e*5 molecules/cell). Likewise, the estimate of viable range 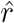 might be made with a model that predicts good enzymatic function at expression levels within a range of 2*e*3 to 5*e*4 molecules/cell, with an empirical accuracy *m* of 3-fold standard deviation on either bound (i.e., 95% chance that the true range of viability has a lower bound between 6.7*e*2 and 6.0*e*3 and an upper bound between 1.7*e*4 and 1.5*e*5, assuming that *m* and *μ* are themselves well known). In this case, the promoter would produce good enzymatic function if the true values turn out to be close to the estimated value but fail if they are farther away. For example, the system will fail if the measurement produced an underestimate, with the true expression level υ being 8.0*e*4 molecules/cell and the model produced an overestimate, with the true upper bound of *r* being 3.0*e*4 molecules/cell.

The probability of success *p*_*s*_ of an engineered design can then be conservatively bounded as being no greater than the probability that a system that has been estimated to have a value in the viable range actually does in fact have that value in the viable range:

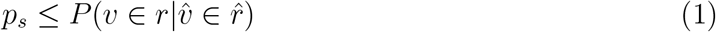

If we evaluate Equation 1 in this context, then by following the derivation provided in the Methods we may compute a conservative bound on the probability of success *p*_*s*_, which we term the *Engineering Error Inequality*:

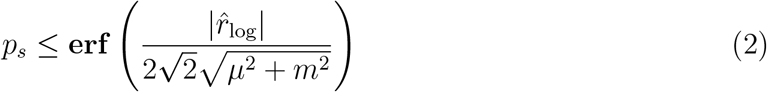

where *μ* and *m* are the geometric standard deviation (log fold) of measurement and model error distributions, respectively,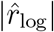 is the magnitude of 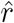 on the log scale (i.e.,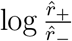), and **erf** is the standard error function. The Engineering Error Inequality thus provides a bound on the probability that an instantiation of a design satisfies its specification, establishing a four-way relationship between the predictive accuracy *m* of a model, the precision *μ* of property value measurements, the estimated range 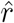 over which a property value will fulfill its specification, and the probability *p*_*s*_ that a particular implementation will actually fulfill said specification.

### Estimating Measurement Precision Requirements

Figure 1(b) illustrates how the Engineering Error Inequality may be thought of with respect to a total “budget” for error in both modeling and measurement. For example, consider a potential definition of “significant” chance of failure (*p*_*err*_ = 1 − *p*_*s*_ = 0.05). The transition around this range is fairly sharp: small changes in total measurement and model errors result in large changes in the likelihood that the system operates as intended. A *p*_*err*_ = 0.05 cutoff is approximately equivalent to two geometric standard deviations under the assumptions of Equation 2, so avoiding a significant chance of failure requires the geometric standard deviation of total error be less than the fourth root of the target range (i.e., the range being two geometric standard deviations up and two down from its geometric center). In current engineering of genetic regulator networks, the estimated range of values 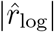 that satisfy a design specification typically spans one or two orders of magnitude (e.g., the range of gene expression levels for a repressor that are high enough to fully repress the promoter it targets, but not too high for the cell to sustain). At these levels, selecting a design that places a value within a 10-fold range requires that error 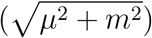 be less than 1.78-fold while doing the same for a 100-fold range requires that error be less than 3.16-fold.

So far, we have discussed the combined error of both model and measurement. Equation 2 tells us that our ability to engineer effectively will be limited by the larger of the two errors. Improvements in the smaller error can improve the situation by a ratio of up to 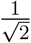 (i.e., about a third). Thus, improving one of the two errors can make a significant difference, but the situation rapidly becomes one of diminishing returns.

Of the two, model uncertainty *m* is generally the more difficult to quantify and control in novel contexts, given its relationship to fundamental unknowns about biology. For example, a prediction regarding transcriptional activity may be thrown off by an unexpected contextual sequence interaction out of scope of the model that is being applied. The precision of measurements *μ*, on the other hand, can be estimated through interlaboratory studies. Even though our understanding of the implications of a measurement is imperfect, we can at least determine how precisely that measurement can be reproduced. Focusing specifically on the question of requirements for measurement precision *μ*, Figure 1(c-e) illustrates this relationship by evaluating Equation 2 to determine the minimum measurement error *μ* for which a desired probability of failure *p*_*err*_ can be achieved given a specified target range size 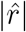 and model error standard deviation *m*. In particular, Figure 1(c) shows a range of 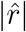 and *p*_*err*_ given a model error with standard deviation *m* = 1.2-fold, Figure 1(d) shows the same for *m* = 1.5-fold, and Figure 1(e) for *m* = 2.0-fold. This range of values is typical for the observed levels of accuracy demonstrated by current predictive models for various aspects of gene expression networks^2–8^. In order to achieve a 5% chance of failure on a 10-fold target range in the first case, with *m* = 1.2, measurement precision needs only to be approximately 1.7-fold, while with *m* = 1.5 the required precision tightens to approximately 1.5-fold, and with *m* = 2.0 the desired chance of failure cannot be achieved even with perfect measurements (*μ* = 0).

Complementarily, more tolerant system designs and better-performing devices can relax demands for both measurement precision *μ* and model accuracy *m* by providing a larger acceptable range of values to target. For example, with *m* = 1.5-fold model accuracy, changing from a 10-fold target 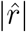 to a 30-fold or 100-fold target 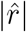 decreases the required measurement precision *μ* to 2.16-fold and 3.01-fold, respectively.

These analyses are conservative, meaning that in practice the *de facto* requirements for effective engineering may be significantly more stringent. However, these estimates can at least be used to determine minimum requirements for use of a measurement technique in engineering biological organisms. Thus, given the typical prediction power of current models and the typical operating ranges of current biological devices^6,8,10,14–20^, we may set a nominal 1.5-fold standard deviation target for measurement precision *μ* as an approximate mid-point in the range where 5% change of failure is achieved with 1.2-fold to 2.0-fold *m* models and 10-fold to 100-fold 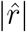 devices, per Figure 1(c-e). Further improvements will bring additional marginal gains, with rapidly diminishing returns for anything tighter than approximately 1.2-fold. On the other hand, measurement precision worse than approximately 2.0-fold is effectively unusable for predicting the behavior of anything but the most tolerant systems and highest performance devices.

Given the landscape defined by evaluating Equation 2 against the current state of the art in modeling and available devices, we suggest that these numbers provide a reasonable set of first approximation targets against which to evaluate modalities of measurement. This, then, is the first key result of this work: in order for a measurement to support effective engineering of genetic regulatory networks given current model precision and device characteristics, we recommend a target precision *μ* of 1.5-fold standard deviation, with a maximum allowable value of 2.0-fold and no significant improvement expected for *μ* less than 1.2-fold.

### Precision of Gene Expression Quantification with Flow Cytometry

We now apply these results to ask whether flow cytometry can satisfy the requirements for measurement precision. Flow cytometry is one of the most frequently used methods for quantifying gene expression, e.g., by co-expressing a fluorescent protein with a gene of interest or by using dye-tagged antibodies that attach to membrane proteins. These instruments produce data in arbitrary units whose values depend on the specific instrument and its configuration, but these data can be mapped to reproducible units by measuring a calibrant of known fluorescence and scaling the unknown data accordingly. These units are given as Molecules of Equivalent Soluble Fluorophores (MESF) or MEx where x is a particular fluorophore, e.g., MEFL for Molecules of Equivalent Fluorescein. Flow cytometry is also the only currently widely accessible assay modality that can collect expression data from large numbers of single cells, which can make flow cytometry particularly valuable for modeling and prediction. Indeed, flow cytometry has been the primary assay used in a number of recent studies demonstrating effective prediction of multi-element genetic expression systems from models of their constituent components^4–6–8–10^. Alternatives either gather population data (e.g., plate reader, RNAseq, proteomics) or single cell data from relatively small numbers of cells (e.g., microscopy) or are not yet widely accessible (e.g., droplet microfluidic sequencing^23^, various single-cell transcriptomic methods^24^).

Prior results with calibrated flow cytometry suggest that this method is indeed likely to be able to achieve the required degree of precision. A study by the International Society for Advancement of Cytometry (ISAC) and the US National Institute of Standards and Technology (NIST) demonstrated that currently available fluorescence calibration beads achieve no better than approximately 1.2-fold variation^25^, a number consistent with prior studies on the variation of dye-based quantification of fluorescence (e.g., ^26^). This provides an anticipated lower bound on measurement error, since measurements that are calibrated using fluorescent beads cannot be more precise that the measurement of the calibrant. Additionally, two interlaboratory studies organized by iGEM^11,12^ suggest a likely upper bound. In these studies, each laboratory independently transformed several genetic constructs into cells, then cultured the cells and measured their fluorescence using plate readers and/or flow cytometry. The flow cytometry measurements in these two studies, with units determined using SpheroTech calibration beads^27^, produced values with geometric standard deviations of 1.7-fold^11^ and 2.3-fold^12^, but they include possible error introduced by independent transformation and culturing methods. To the best of our knowledge, no prior study has specifically examined the precision of measurements quantifying fluorescent protein expression via bead-calibrated flow cytometry.

### Study Design and Instruments

To address this question, we organized an interlaboratory study to measure the same cells on different instruments. Figure 2 shows the overall design of this study. First, five strains of *E. coli* were transformed and cultured in a single laboratory, each strain with a different plasmid: a blank plasmid expressing no fluorescent protein, two plasmids for constitutive expression of GFP at different expression levels, and two plasmids for constitutive expression of mCherry at different expression levels (full details are provided in S1: Interlab Plasmid Designs). All participating laboratories were sent frozen aliquots of each of the samples, along with SpheroTech RCP-30-5A rainbow calibration beads^27^, such that all measurements were made with aliquots from the same biological material and the same set of calibration beads. The laboratories performed flow cytometry measurements in two stages, first using one set of samples to determine appropriate instrument settings, then measuring three sets of samples to acquire study data. For instruments with variable channel gain, each sample was measured at three different gain levels. Three technical replicates of measurement were requested for each sample in every measurement condition. All data was then uploaded for a unified analysis process. Full details are provided in Methods, S2: Protocol, and S4: Flow Cytometry Data Processing.

**Figure 2:**
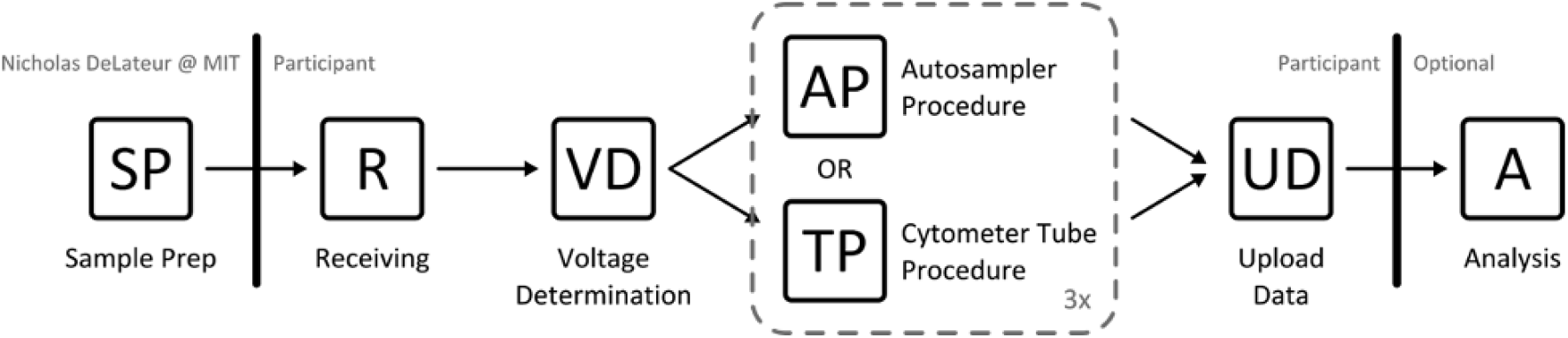
Interlaboratory study design: samples prepared in one lab were split and shipped to all participant labs. Once received at each lab, samples were measured first to determine instrument settings, then to acquire study data, which was uploaded for unified analysis.

In total, measurements and configuration information were collected from 23 flow cytometers at 15 institutions. We deliberately did not attempt to select for identical instruments, but instead gathered as much information as was available about each instrument in order to allow our study to consider both identical and non-identical channels and instruments. The full list of instruments and channels is provided in S3: Participating Instruments and Channels. In total, the study comprised 12 different models of flow cytometer from three manufacturers. The overall variety of available channels is quite wide. For subsequent analysis, we selected channels that best matched the excitation and emission spectra of GFP and mCherry, as shown in Figure 3 (details given in S4: Flow Cytometry Data Processing). For GFP, the 488nm excitation, 530nm/30nm emission filter channel is widespread (16 of 23 instruments), with all but two others closely similar. For mCherry, channels were more heterogeneous, with the most frequent being the 561nm excitation, 610nm/20nm emission filter channel (9 of 23 instruments).

**Figure 3:**
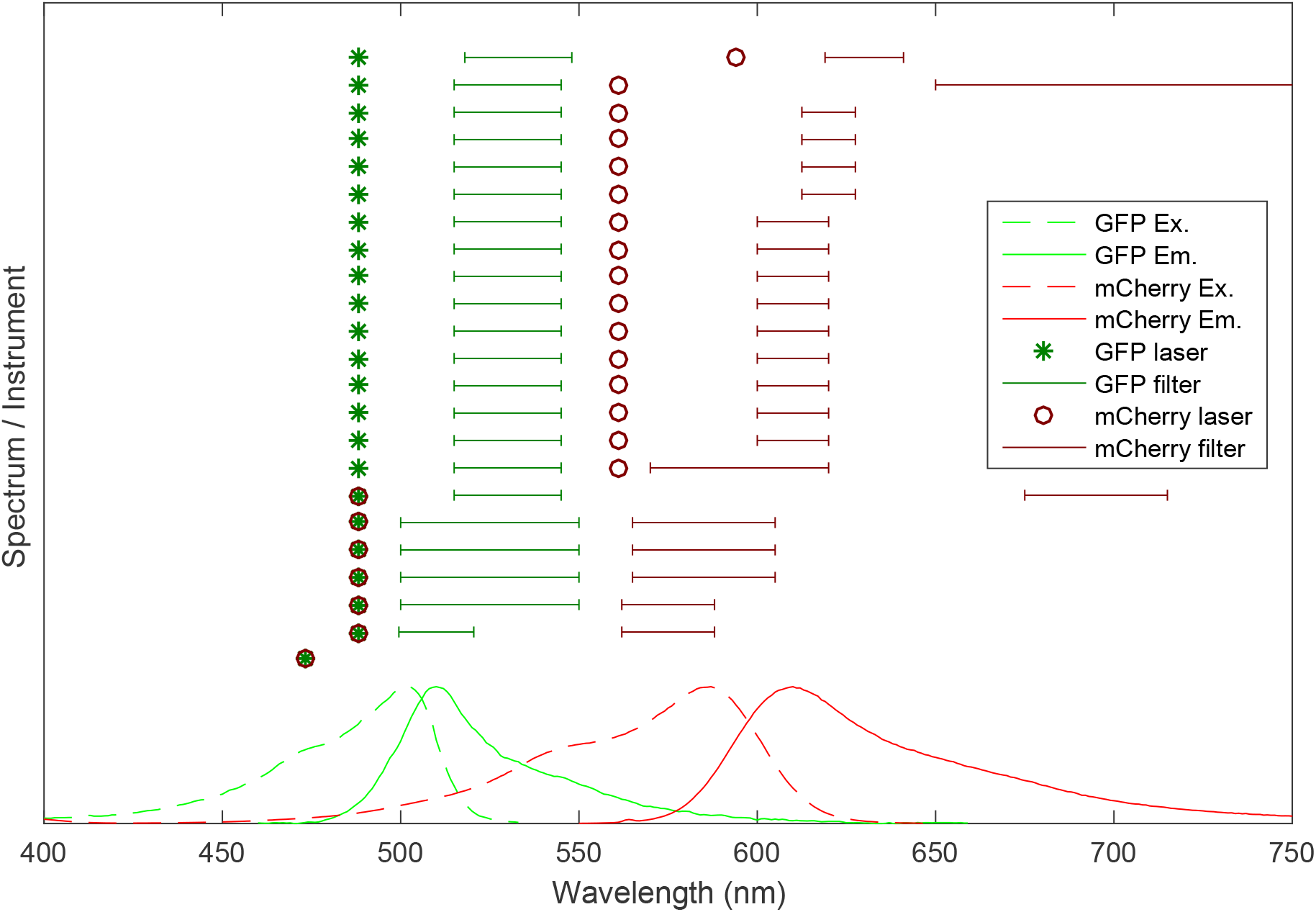
Comparison of best match channels for calibration of GFP to MEFL and and mCherry to MEPE using SpheroTech RCP-30-5A beads (sorted first by laser then by emission filter center). Relative spectra for GFP and mCherry are plotted in the background for comparison (source: fpbase.org). For GFP, most channels are quite similar, while for mCherry there is a high degree of heterogeneity. Note: for the 473 nm excitation instrument at bottom, emission filter information was not available and is thus not shown. Full channel details information is provided in S3: Participating Instruments and Channels.

In principle, every set of channels with identical excitation and emission filters should produce identical results. For non-matched channels, the degree of disparity should depend on the relative difference between the spectral properties of the calibration beads versus the spectral properties of the fluorescent proteins. In principle, such spectral information might be used to adjust unit to be equivalent across unmatched channels. For the present, however, we will ask only the empirical question of how much variability is observed for both matched and unmatched channels.

Furthermore, in all cases, we cannot truly say that the calibration is to the designated units of the manufacturer (molecules of equivalent fluorescein, MEFL and molecules of equivalent phycoerythrin, MEPE), since no channel precisely matches the channels for which the calibrant manufacturer has provided values. As the target of our study is not absolute units but variation in measurement, however, this is not a limitation. We will use the designation nonetheless as a shorthand for beads calibrated using the values provided for MEFL and MEPE in the closest matching available channel. Note that this lack does indicate a need for coordination between calibration bead manufacturers and flow cytometry providers on standard channels and/or more complete spectral information to be made available about calibration beads.

### Study Results

Of the 23 participating instruments, 19 had adjustable gain settings (enabling measurement at 3 sensitivities) and 4 had fixed gain, meaning that overall 61 data sets were collected. Each should have 10 technical replicate groups (5 strains measured on 2 fluorescence channels) for a total of 610 anticipated technical replicate groups, with three replicates per group. Data was analyzed following the process in S4: Flow Cytometry Data Processing, with geometric statistics computed for the cell-like events of each sample (geometric statistics are used in all cases due to the expected log-normal distribution of strong gene expression^21^). Instrument and sample errors, however, meant that not all groups had three replicates and not all technical replicate groups could used for analysis. On one instrument, only one replicate was able to be collected and on another the selected red channel was found to be inoperative, invalidating 40 technical replicate groups. Another 86 replicate groups were deemed invalid due to insufficient data, having fewer than two technical replicates with at least 100 cell-like events per replicate. This left a total of 484 replicate groups accepted for analysis.

Variation between technical replicates can provide an empirical baseline for evaluating instrument precision. At the same time, once a baseline has been established, deviations from that baseline are likely to indicate issues in protocol execution that are cause for data exclusion. Figure 4(a) shows the distribution of standard deviations of means across the replicate groups. Overall, technical replicates show a consistently high level of precision, with more than half of all replicate groups having a standard deviation of less than 1.04-fold and 84.3% having a standard deviation of less than 1.2-fold. Above approximately 1.2-fold, however, standard deviations increase sharply. Both the low-event samples and the high-variation replicate groups are distributed across instruments and intermixed with other conditions that had no issues, indicating problems are likely due to issues with particular samples or executions rather than systematic in nature. We thus exclude all high-variation technical replicate sets from the remaining comparisons across voltage levels and instruments, using a threshold of 1.2-fold standard deviation, due to the likelihood that these samples would distort results with error unrelated to the subject of study. Hereafter, we will use *accepted replicate group* to refer to technical replicate groups that are both valid and have less than 1.2-fold standard deviation.

**Figure 4:**
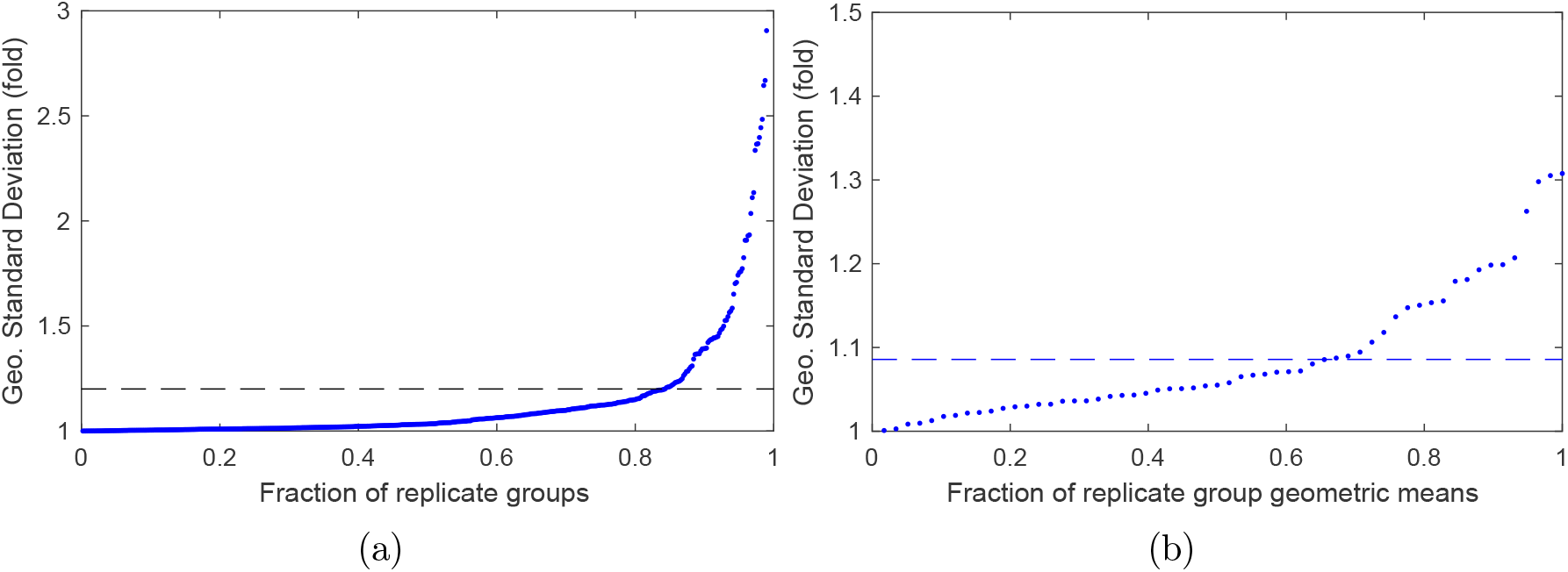
(a) Distribution of geometric standard deviation of means over replicates, across a total of 484 replicate groups. The dashed line shows a cut-off threshold of 1.2-fold. (b) Distribution of geometric standard deviation of replicate group geometric means for fluorescent samples measured at different voltages on the same instrument, across a total of 58 sets of voltage-varying replicate groups. The dashed line shows the geometric mean of geometric standard deviations, which is 1.086-fold.

We then turned our attention to comparisons of the values obtained from different configurations of a single instrument or from different instruments. On an individual instrument, calibration beads should map data obtained with different channel voltage settings to identical units as long as the measurements are not saturated at the high or low end of the quantifiable scale. We thus compared the green fluorescence measurements for GFP-expressing strains and red fluorescence measurements for mCherry-expressing strains, omitting the non-fluorescent strain and channels. For these conditions, we considered all sets of accepted replicate groups in where there are multiple groups identical except for channel voltage settings (i.e., same sample, same channel, and same machine). There are a total of 58 such sets, and Figure 4(b) shows the geometric standard deviation of the replicate group geometric means for each of these sets. Overall, we confirm the expected result, that calibration beads generally allow close comparison across voltage levels, with the geometric mean of geometric standard deviations equal to 1.086-fold and even the maximum observed geometric standard deviation equal to only 1.31-fold.

To compare fluorescent sample measurements between instruments, we restricted the comparison to one gain level for each fluorescent sample and selected the technical replicate group with the lowest standard deviation. In theory, channels with identical excitation and emission parameters should produce identical values once calibrated, even though they are on different instruments. Figure 5 shows the geometric standard deviation for each fluorescent sample for each cluster of instruments with matching channels. Overall, the differences are notably higher than for different voltages on a single instrument. The geometric mean of these geometric standard deviations is still only 1.52-fold. Notably the only two channels shared by high numbers of instruments (488 ex, 530/30em for GFP; 561 ex, 610/20 em for mCherry) have values very close to this for both the high and medium constructs. These results indicate that flow cytometry with bead-based calibration can meet the 1.5-fold target precision estimated above.

**Figure 5:**
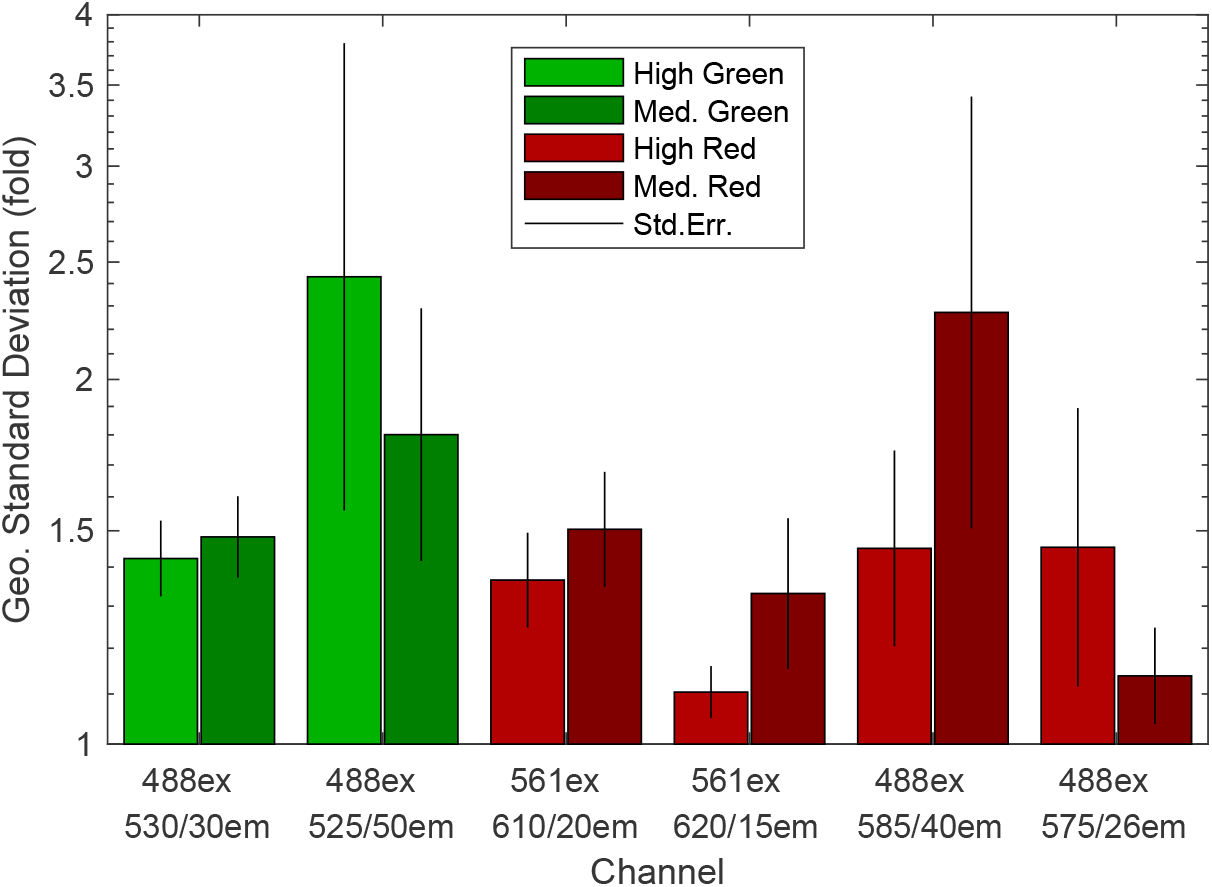
Comparability of fluorescence measurements across instruments with equivalent channels. Bars show the geometric standard deviation of the lowest standard deviation technical replicate group of fluorescent samples per instrument for groups of instruments with channels having identical excitation and emission filter values, along with the standard error for this estimate. Bar color indicates GFP or mCherry, with the high expression construct on the left and medium expression construct on the right of each pair of bars. The geometric mean of geometric standard deviations is 1.52-fold.

Finally, we consider the relationship of calibrated values as measured by instruments with different channels. Figure 6 shows these values for each of the four fluorescent constructs. For GFP, three channels (488 ex, 530/30 em; 488 ex, 525/50 em; 488 ex, 533/30 em) produce values for which the variation between channels is less than the variation within channels (though one of the 525/50 instruments provides a potentially concerning outlier). The 488 ex 510/21em channel, appears marginal (though this is uncertain given there is only a single data point) and the 473 ex channel with unknown emission filter is clearly registering outside the range of any other channel. For mCherry, there are also three channels with lower variation between channels than within channels (561 ex, 610/20 em; 561 ex, 620/15 em; 594 ex, 630/22 em). The others are all distinct from this set of three, but there are too few instruments with these channels to determine whether 488 ex, 695/40em matches 561 ex, 650LP em or whether 488 ex, 585/40 em matches any of the three other channels with which its range of values overlaps.

**Figure 6:**
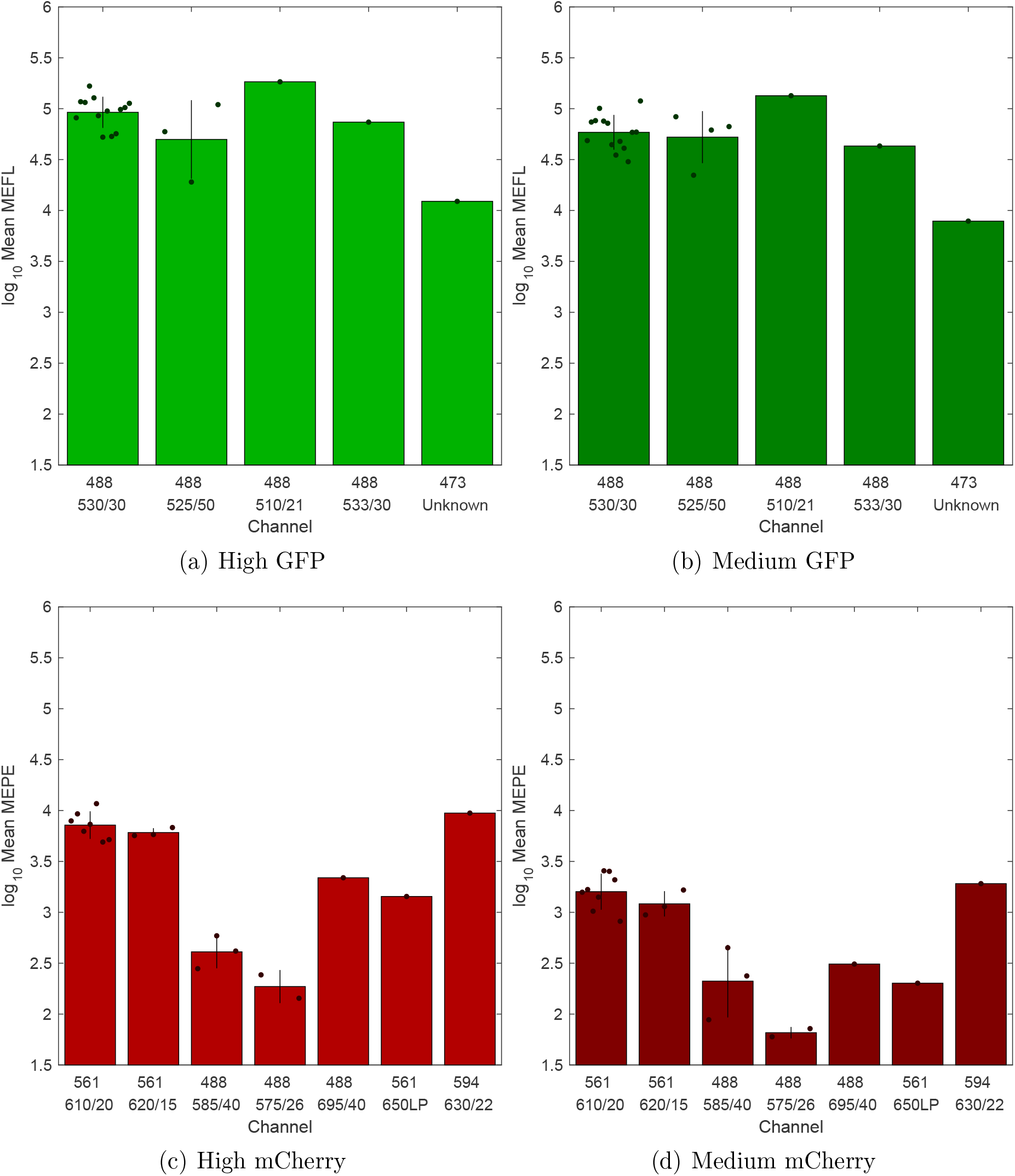
Comparison of calibrated values measured by instruments with different channels for each of the four fluorescent constructs. Dots show the geometric means of the lowest standard deviation technical replicate group for each instrument, bars show the geometric mean and standard deviation over groups. For channels shared by more than one instrument, the vertical bar shows the geometric standard deviation. Overall, channels with similar emission filters appear to produce sufficiently comparable results, despite not being optically matched.

Overall, this indicates that it is possible for channels with similar emission filters to produce results that are sufficiently comparable for effective engineering, despite an imperfect optical match. In particular, this is supported by the similarity of distributions observed for the 488 ex, 530/30 em and 488 ex, 525/50 em channels for GFP and for the 561 ex, 610/20 em and 561 ex, 620/15 em channels for mCherry. Channels with a higher degree of difference, however, clearly produce different values, as demonstrated, for example, by the comparison of the 561 ex, 610/20 em and 488 ex 575/26 channels for mCherry. More investigation is needed, however, to determine the bounds of tolerable variation between channels, the degree to which other requirements can also be relaxed (e.g., using calibrants from different lots or different manufacturers), and the degree to which more sophisticated unit computation can compensate for differences between channels.

## Discussion

The results that we have presented establish an error budget for engineering genetic regulatory networks, with the *Engineering Error Inequality* quantifying the tension between device characteristics, error in measurement and modeling, and the difficulty of engineering. Once established, this relationship can be used to derive requirements for any one of these properties based on the available capabilities for the others. In particular, with the current state of the art in genetic regulatory network engineering, we have estimated a target precision for measurements of *μ* = 1.5-fold geometric standard deviation, with approximate upper and lower bounds of 2.0-fold and 1.2-fold.

By measuring a uniform set of cell samples on a heterogeneous collection of flow cytometers, we determined that bead-calibrated flow cytometry can indeed meet this error budget, with a geometric mean value of 1.52-fold geometric standard deviation across matched channels. At this level of precision, we should expect that flow cytometry can be effectively used for engineering with reasonably good models (*m* ≤ 2.0-fold), using biological devices with strongly differentiated outputs 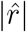 at least 10-fold to 100-fold, depending on *μ* and *m*), collectively meeting a sufficiently tight bound on predictability. Flow cytometry measurements do not, however, appear to be sufficiently precise to allow for engineering with either poor models or weak devices. Bead-based calibration is also sufficient to allow comparison of data from channels with similar emission filters, even when these filters are not matched. As a byproduct of this study, we also confirm that bead-based calibration allows highly precise matching across different voltage levels on a single instrument (finding a precision of 1.09-fold), which has the potential to expand effective measurement range and to allow fusion of otherwise incompatible datasets.

Based on these results, we recommend that the following be adopted as best practices for use of flow cytometry measurements as a basis for engineering genetic regulatory networks:

- Calibrate units using independent reference materials (e.g., NIST traceable dye-based calibration beads). Use of a reference strain (e.g., RPU^6,28^) is not sufficient, as demonstrated in^11^.
- Collect technical replicates for controls in triplicate to validate that measurement replicability on the instrument is better than 1.2-fold standard deviation.
- Use negative and single positive controls for background removal and compensation (not part of our study, but standard recommended practice, per^29,30^)
- Standardizing machine, light path, and laser power are not necessary, as long as calibration beads are used and controls indicate the instrument is performing well.
- Standardize laser/filter combinations as much as possible, even at the cost of somewhat less precise match with fluorescent protein properties. Based on results here and elsewhere, we recommend the use of the following channels for any fluorescent protein with a reasonable optical match:

– Yellow/green fluorescence: 488nm excitation, 530/30 emission (with 525/50 an acceptable alternative)
– Red fluorescence: 561nm excitation, 610/20 emission (with 620/15 an acceptable alternative)
– Blue fluorescence: 405nm excitation, 450/50 emission^1^

The final recommendation also points to important additional work that is still needed. Notably, coordination between calibration bead manufacturers and flow cytometry providers to standardize channels and/or to provide more complete spectral information about calibration beads would allow for device characterization data to be shared across a broader range of laboratories and instruments. Likewise, with a profusion of available fluorescent proteins, it would benefit the engineering community to consider reusability of data when selecting fluorescent proteins. There is also a gap between calibration in terms of equivalent dye molecules and the ability to interpret these numbers in terms of the actual fluorescent protein used, as illustrated by the high degree of variation between channels in Figure 6(c-d). Shifting units from equivalent dye molecules to protein molecules would also address the problem of channel-to-channel conversion, though in some circumstances this can also be handled by multi-color controls, as shown in^5,31^.

Beyond flow cytometry, it will be important establish that other measurement modalities can also achieve the requisite level of measurement precision. Plate reader data seems readily able to achieve similarly usable, if imperfect, error levels, given the precision and unit equivalence demonstrated in^12^, though this has not yet been explicitly confirmed. Calibration of RNAseq with spike-in controls also appears likely to provide sufficient precision, based on the results in^13^. Other forms of measurement likewise may be evaluated using similar methods to those presented here.

Finally, effective engineering will only be actually enabled by the availability of sufficient data collected using methods such as these. The synthetic biology community has built up large public repositories of reusable genetic parts, through organizations such as iGEM, Addgene, and SEVA. The ability to engineer with such parts will be advanced by the community building up public repositories of reusable data describing part behavior, with the data collected using comparable fluorescent proteins and sufficient precision and calibration to allow its application in the design of new genetic regulatory networks.

## Methods

### Derivation of Engineering Error Inequality

Consider an engineered design with a collection of measurable property values υ, such that the design will meet its specification only if every property value is within some interval range *r* = [*r*_−_, *r*_+_] specified for that property.

Note that this is formulated as a necessary condition, but not sufficient: there may be other interactions within the system that impose additional constraints on relationships between properties, but we may at least take the conservative view that success cannot be achieved without at least getting every property to a value within that property’s specified range of potentially acceptable values. Likewise, we may assume without loss of generality that there is only a single range of viable values: a disjoint range *r* = [*r*_−,1_, *r*_+,1_] ∪ [*r*_−,2_, *r*_+,2_] is strictly lower probability than a single range of the same total magnitude *r*′ = [*r*_−,1_, *r*_+,1_ + (*r*_+,2_ − *r*_−,2_)], as the impact of uncertainty is greatest at an interval boundary and a disjoint range has more such boundaries.

The probability of a particular instantiation of a design satisfying its specification is thus bounded above by the joint probability of all property values being within their respective ranges. This, in turn, is bounded above by the probability of any individual value being within a specified range, such that *p*_*s*_, the probability of successful realization of a design specification with a particular set of design choices, given an estimated property value 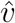 whose distribution is determined by measurement precision, and an estimated range 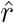 whose accuracy is limited by the accuracy of available models, may be bounded above by:

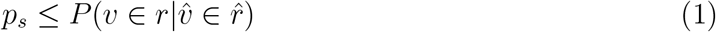

where *s* is the event of success, in which an instantiation of a design is able to meet its specification.

The distribution of genetic expression levels in cells is typically log-normal^21^ (or a close approximation^22^), which implies that estimates of expression levels made from measurements will typically have a log-normal estimation error distribution as well, unless another larger source of error dominates (e.g., stochasticity due to very low expression levels). Likewise, effective quantitative predictive models have typically shown distributions well-described in terms of error on a logarithmic scale as well (e.g.,^3–8^). When dealing with genetic regulatory networks, then, a reasonable null hypothesis is that both measurement errors and model errors have independent log-normal distributions.

When this hypothesis holds, the composition of these two perturbations is a log-normal with the sum of their variances:

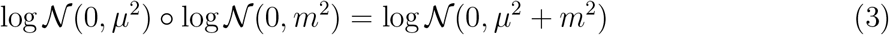

taking *μ* as the (log-scale) standard deviation of the distribution of measurement error and *m* likewise for model error.

If it is possible to choose any value for 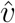, then *p*_*s*_ may be maximized by choosing 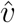 to be at the center of 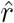 (if not all values can be chosen, the best available value may have an even lower chance of success, so the inequality is unaffected). The actual values of 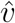 and 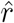 then become irrelevant, and we may refactor the equation in terms of the magnitude of the estimated range versus the magnitude of uncertainty from model and measurement around a zero mean:

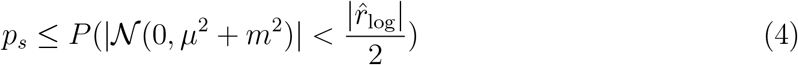

with the magnitude of the range in the equation is also quantified on a log-scale (note however, that this equation does assume that the values of *μ* and *m* are well constrained).

We can then normalize the range to express this probability in terms of the number of standard deviations with respect to a standard normal distribution:

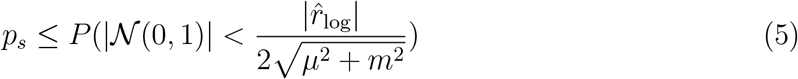

This value of this probability is the cumulative distribution for a folded normal distribution evaluated at 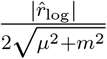, which is:

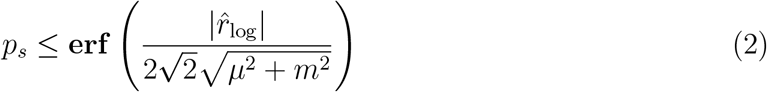

where **erf** is the Gauss error function. We thus have an inequality that relates model accuracy, measurement precision, and design tolerance (in terms of magnitude of acceptable range) to the probability that an engineered genetic regulatory network satisfies its design specification.

### Plasmid construction

Plasmids were constructed by Golden Gate^32^ assembly in the MoClo^33^ architecture.

### Strain Culturing

Plasmids were transformed into 10-beta competent *E. coli* cells (New England Biolabs cat. #C3019H) according to the manufacture’s recommendation. Single colonies were struck out on selective media and then grown overnight in 100 μL of M9 minimal media supplemented with glycerol^34^ and carbenicillin (100 μg mL^−1^) at 900 RPM and 37 °C (Model DTS4, ELMI North America). After 16 hours each strain was diluted 15 μL into 185 μL of media twice successively and grown at 900 RPM and 37 °C. After 3 hours each strain was diluted 15 μL into 185 μL of media twice successively and grown at 900 RPM and 37 °C for another 5 hours. At this point all strains had an OD600 of 0.5 or less and were concentrated to an OD600 of 0.5; 150 μL of this culture was combined with 90 μL of 0.45 μm filter-sterilized 50% glycerol and mixed gently, then kept on ice. Pre-labelled 2 mL labelled Eppendorf tubes chilled to −80 °C were then used to aliquot 10 μL of culture and immediately placed back at −80 °C until distribution.

### Flow Cytometry

#### Determining Instrument Pre-Collection Gating

Each recipient was instructed to set FSC and SSC voltages (if permitted by their instrument) such that the peak density of cells was centered in the range of their detector. As pre-collection gating options can vary significantly by instrument and software, recipients were instructed to set gating to retain as many cell events as possible, even if this would result in more non-cell events being captured in the same. Additional details can be found in S2: Protocol. See Data Processing and S4: Flow Cytometry Data Processing for the separation of cell and non-cell events post-capture.

#### Determining High, Medium, and Low Channel Voltages

Each recipient was instructed to take the brightest strain for red and green and raise the corresponding channel’s voltage until the distribution was as high as possible without clipping the sample. This would be used as the High voltage; the process was repeated with targeting voltages for 10x and 100x fold less arbitrary units for Medium and Low voltage settings respectively. For instruments with low maximum values, recipients were instructed to still space the Medium and Low voltages evenly, but to lower the overall separation so that the sample was not clipped significantly on the low end. For instruments without a variable voltage setting, this step was necessarily omitted. Additional details can be found in S2: Protocol.

#### Sample Preparation

Bacterial cultures were prepared for flow cytometry as follows: Remove one replicate of samples from −80 °C and transfer to 42 °C heat bath for 60 sec. 990 μL of sterile filtered 1X PBS to bacterial culture, or 75 μL for calibration beads, is added to bring the sample concentration to 1X for running on the flow cytometer. Samples are then run immediately. This procedure is repeated independently for each replicate.

#### Event Collection

Samples were run as close to these conditions as possible based on local instrument and software: Each sample 1 μL s^−1^ for either 150 μL or 10^5^ total events, whichever was achieved first.

## Supporting information

S1: Interlab Plasmid Designs

S2: Protocol

S3: Participating Instruments and Channels

S4: Flow Cytometry Data Processing

S5: Sample Statistics

## Data processing

Cell events were then separated from non-cell events by means of a Gaussian mixture model filter trained on the negative sample. Flow cytometry data was processed using the TASBE Flow Analytics software package^31^, using the recommended practices for gating, background subtraction, and bead-based calibration. Additional details and examples are provided in S4: Flow Cytometry Data Processing.

## Author Contributions

- Conceptualization: J.B., B.T. J.S. S.C.-H., N.A.D.
- Data curation: J.B., B.T., J.S., S.C.-H.
- Formal analysis: J.B., J.S.
- Investigation: Experimental data gathered by Calibrated Flow Cytometry Study Contributors, plus N.A.D., J.S., S.C.-H. (full list in Acknowledgements section)
- Methodology: J.B., B.T. J.S. S.C.-H., N.A.D., M.S.
- Project administration: J.B., B.T. J.S. S.C.-H., N.A.D.
- Funding acquisition: J.B., J.J.T., R.W.
- Writing (original draft): J.B.
- Writing (review & editing): J.B. (anyone who responds)

## Acknowledgement

Partial support for this work was provided by NSF Expeditions in Computing Program Award #1522074 as part of the Living Computing Project.

This document does not contain technology or technical data controlled under either the U.S. International Traffic in Arms Regulations or the U.S. Export Administration Regulations.

Consortium members (sorted by institution name) collected data:

- Leidy D. Caraballo, Augusta University, Augusta, GA, USA
- Joel M Sederstrom, Baylor College of Medicine, Houston, TX, USA
- Wilson Wong, Boston University, Boston, MA, USA
- Karmella A. Haynes, Emory University, Atlanta, GA, USA
- Alina Chan, Harvard Medical School, Boston, MA, USA
- Leo d’Espaux, Jasper Therapeutics, Redwood City, CA, USA
- Maren Wehrs, Joint BioEnergy Institute, Emeryville, CA, USA
- Claire K. Sanders and Taraka Dale, Los Alamos National Laboratory, Los Alamos, NM, USA
- Karen Dwyer, UT MD Anderson Cancer Center, Houston, TX, USA
- David Ross, National Institute of Standards and Technology, Gaithersburg, MD, USA
- Alexandra Westbrook and Julius Lucks, Northwestern University, Evanston, IL, USA
- Alon R. Azares, Texas Heart Institute, Houston, TX, USA
- Nicolas Loof, University of Texas Southwestern, Dallas, TX, USA
- Joo-Young Lee, Kyung Kim, Bryan Bartley, and Herbert Sauro, University of Washington, Seattle, WA, USA

## Supporting Information Available

### S1: Interlab Plasmid Designs

GenBank files for the five plasmid designs used in the study.

### S2: Protocol

Protocol provided to laboratories collecting data for the study.

### S3: Participating Instruments and Channels

List of instruments used in study and all information that was able to be collected regarding the available channels for each instrument, highlighting channels selected as “best match” for GFP and RFP measurements.

### S4: Flow Cytometry Data Processing

Details of how flow cytometry data processing was conducted.

### S5: Sample Statistics

Table of statistics for every sample evaluated.

## TOC Graphic

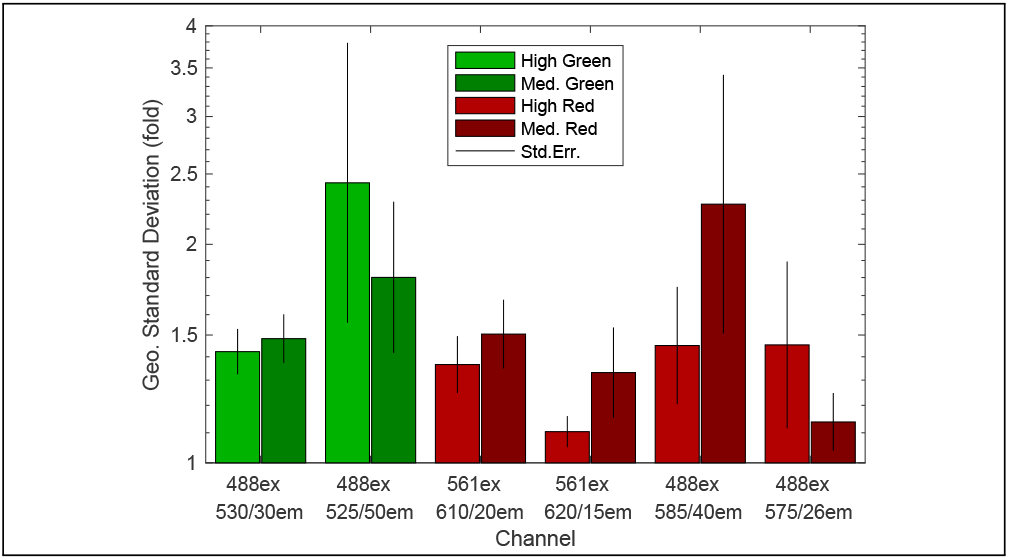

## Footnotes

1 We found this channel to be widespread among cytometers (S3: Participating Instruments and Channels); though blue fluorescence was not analyzed in this study, measurements of this channel have been sufficiently precise to support high-accuracy predictions in prior studies^4,5,7,10^.

