## Supplementary material for "Meeting Measurement Precision Requirements for Effective Engineering of Genetic Regulatory Networks": S2: Protocol

### Hypothesis that we will test:

We propose a method for comparison of fluorescent measurements based on beads. We hypothesize that with this method:

- Repeatability can be achieved with better than 1.5x precision (based on ISAC interlab with beads).
- Reproducibility can be achieved with better than 2.0x precision.

### Experimental Design

#### Materials:

- Stable **biological samples** that can be shipped to every participating laboratory
    - Frozen *E. coli* cells with green = GFPmut3*, red = mCherry
    - Five expressions: blank, medium green, strong green, medium red, strong red.
        - Strains:
            - Tube T05: sFW05 contains pL1f1_EV
            - Tube T07: sFW06 contains pL1f1362 - Strong GFP
            - Tube T06: sFW07 contains pL1f1357 - Medium GFP
            - Tube T09: sFW08 contains pL1f1372 - Strong mCherry
            - Tube T08: sFW09 contains pL1f1285 - Medium mCherry
            - Correction: the medium and strong strains were swapped for both red and green fluorescent proteins. See Tube ID and contained Strain ID above for clarification.
    - Send five identical stable biological samples for each of the 5 strains (25 samples total)
        - A single stable biological sample contains 10uL of sample
        - For each strain, 1 sample is for instrument detector voltage (a.k.a. “gain”) determination, 3 samples are for the experiment, and 1 sample is a spare
- Fluorescent **calibration beads** capable of being directly compared
    - SpheroTech rainbow beads from lots judged equivalent by SpheroTech: SpheroTech RCP-30-5A beads, lots AD04, AE01, AF01, AF02, AH01, AH02, AJ01
    - Five samples containing 75uL of these beads will be shipped in lightproof tubes
- **Flow cytometers** with equivalent laser/filter combinations for GFP and mCherry
    - GFP: 488 laser, 530/30 filter (or closest equivalent)
    - mCherry: 561 laser, 610/20 filter (or closest equivalent)
    - **All detection channels should be turned on** (not just the “best match” channels), and all of area, width, and height should be recorded.
    - If your flow cytometer does not have similar channels, please still measure on all channels anyway: the results will still be valuable for negative results and investigation of inter-channel conversion.

### Experimental Procedures:

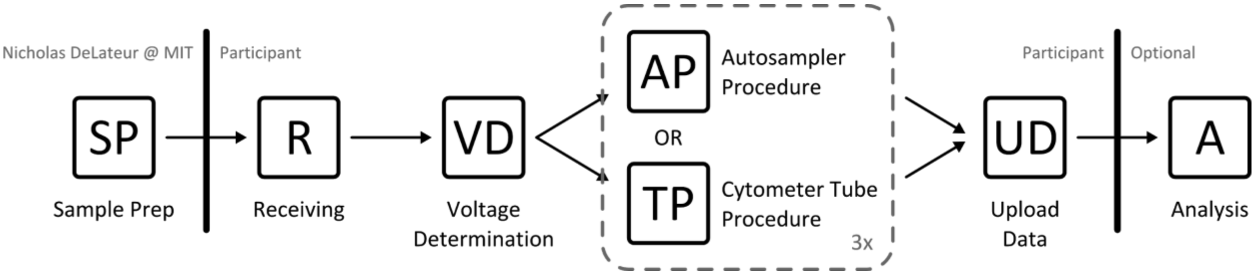


#### SP Stable Biological Sample Preparation Procedure

*This protocol is performed only by the sample-preparing lab (MIT).*

- Streak plate(s) with strain(s) of interest:
    - sFW05 contains pL1f1_EV
    - sFW06 contains pL1f1362 - Strong GFP
    - sFW07 contains pL1f1357 - Medium GFP
    - sFW08 contains pL1f1372 - Strong mCherry
    - sFW09 contains pL1f1285 - Medium mCherry
- Overnight culturing
    - Add 200uL of M9-minimal-glyc + appropriate antibiotics to wells of a 96-well plate
    - Using a small pipette tip, inoculate well with single colony for each strain of interest
    - Seal plate and place in 37C rotating incubator (900 RPM)
- Sub-culturing, round 1
    - Dilute overnight sample into M9-minimal-glyc + appropriate antibiotics via two serial dilutions with a dilution ratio of 3:40 for each dilution
        - On a 96-well plate, add 185uL M9-minimal-glyc + appropriate antibiotics to 2 new wells per strain of interest.
        - Transfer 15uL of well-mixed overnight sample into freshly prepared well. Mix gently.
        - Transfer 15uL of well-mixed diluted sample into freshly prepared well. Mix gently.
    - Seal plate and place in 37C rotating incubator (900 RPM)
- Pre-freeze culturing, round 2
    - After 3 hours of sub-culture in round 1, dilute cultures into M9-minimal-glyc + appropriate antibiotics via two serial dilutions with a dilution ratio of 3:40 for each dilution
        - On a 96-well plate, add 185uL M9-minimal-glyc + appropriate antibiotics to 2 new wells per strain of interest.
        - Transfer 15uL of well-mixed sub-culture round 1 sample into freshly prepared well. Mix gently.
        - Transfer 15uL of well-mixed sub-culture round 1 diluted sample into freshly prepared well. Mix gently.

  - Seal plate and place in 37C rotating incubator (900 RPM)
- Freeze culture
  - After 5 hours of culturing pre-freeze round 2 cultures, transfer 150uL of pre-freeze round 2 culture into new Eppendorf tube.
  - Add 90uL of filter-sterilized 50% glycerol to Eppendorf tube. Mix gently.
  - Aliquot 10uL of glycerol-diluted pre-freeze round 2 culture into new Eppendorf tubes.
  - Place 10uL glycerol-diluted pre-freeze round 2 cultures into -80C freezer.
- Bead sample prep
  - Vortex Spherotech calibration bead dropper bottle gently.
  - Remove top of calibration bead dropper bottle.
  - Aliquot 90uL of calibration beads into lightproof Eppendorf tubes.
  - Place lightproof Eppendorf tubes into 4C storage.

#### R Sample Receiving Procedure:

- Take the samples from the dry ice and put them in -80C storage
- Take the beads from their package and put them in 4C storage

#### VD Voltage Determination Procedure (to be performed once, before first replicate):

- Prepare one PBS-diluted biological sample for each "strong" single-color biological sample, per appropriate procedure below (either Autosampler/Plate Procedure or Cytometer Tube Procedure).
  - Strains for this procedure:
    - Tube T07: sFW06 contains pL1f1362 - Strong GFP
    - Tube T09: sFW08 contains pL1f1372 - Strong mCherry
    - Correction: the medium and strong strains were swapped for both red and green fluorescent proteins. See Tube ID and contained Strain ID above for clarification.
- Tune forward and side scatter voltages so that:
  - FSC and SSC should be set to have the strong cell peak as close to the center of the detector range as possible.

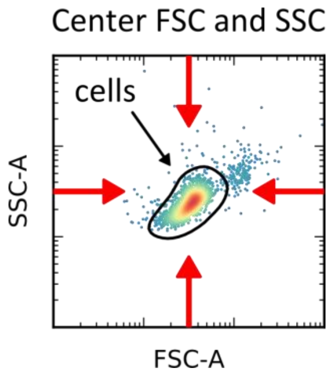

        ■ The FSC and SSC detector voltages for bead samples may be adjusted to ensure that bead events fall within the FSC and SSC detector ranges, but only if necessary (i.e. only if bead events saturate the FSC or SSC channels at the FSC or SSC detector voltages used for measuring biological samples).
    ○ Thresholding in the instrument should be set to ensure that **no cell events are discarded**.

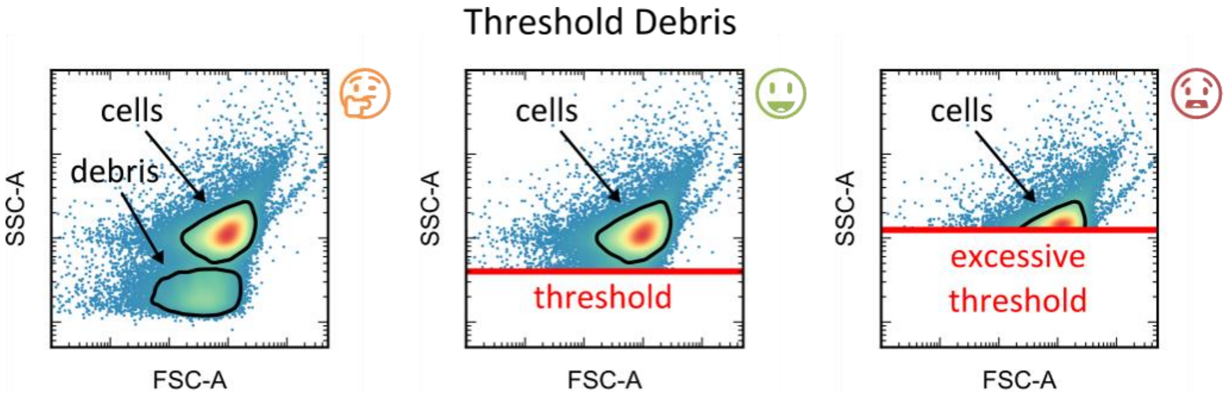


        ■ Examples from past practice:
            ● MIT: 500 SSC, 500 FSC.
            ● NIST Gaithersburg: 200 FSC, 2000 SSC
            ● Rice: SSC, 65%

- For each fluorescent protein (GFP, mCherry), identify channels in which an appreciable nonzero fluorescence signal can be detected (channels with no appreciable fluorescence signal from any fluorescent protein do not need to have their voltages tuned).
    ○ Examples: we expect a strong fluorescence signal from GFP excited by a 488nm laser and measured through a 530/30 filter and from mCherry excited by a 561nm laser and measured through a 610/20 filter. Channels with similar configurations may also detect appreciable amounts of fluorescence, though, and are of interest to us.
- Tune voltages of the identified fluorescence channels:

- For each fluorescent protein / fluorescence channel combination, identify three different voltage settings:

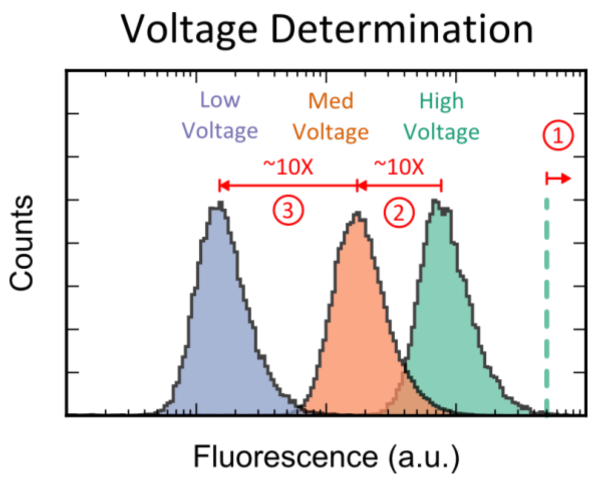


- The goal is to put each high as high as it can safely go without high clipping, then to decrease peak a.u. by ~10x with each lower setting.
  We expect that this will place each high single-color sample at around:
  - 10^6 max: 10^3, 10^4, and 10^5 a.u.
  - 10^5 max: ~ 10^2.5, 10^3.5, 10^4.5 a.u. (as high as can go without clipping)
  - 10^4 max: ~ 10^1, 10^2, 10^3 (as high as can go without clipping)

- Record voltage settings for reuse in replicates.
- If necessary, run a PBS blank sample immediately after a cell sample to document the amount of carryover your cytometer setup experiences. If carryover is detected, run focusing fluid blanks between samples in the below procedures (Autosampler/Plate Procedure or Cytometer Tube Procedure).

#### AP Autosampler/Plate Procedure:

- Prepare >= 20mL of 1X PBS filtered (0.2um). Confirm and record pH of the PBS that is used.
- Transfer 200uL of 1X PBS to each of three wells on plate (one well for each voltage).
- Prepare PBS-diluted samples.
  - Retrieve 1 Eppendorf tube of each strain from the -80C storage and 1 Eppendorf tube of calibration beads from the 4C storage.
  - Immediately place Eppendorf tubes containing strains in 42 degree water bath or heat block for exactly 60 seconds (do *not* place Eppendorf tube containing beads in water bath or heat block).
  - Add 990uL of 1X PBS to Eppendorf tubes containing 10uL of biological sample or 75uL of calibration beads. Vortex gently.
  - Transfer 200uL of each PBS-diluted sample to each of three wells on plate (one well for each voltage) (600uL total of the 1mL PBS-diluted biological samples or 1.08mL PBS-diluted calibration beads sample).

- Order of PBS-diluted samples on plate:
  - PBS, empty vector (EV), medium green (MG), strong green (SG), medium red (MR), strong red (SR), beads
  - Place a focusing/sheath fluid blank (FF) before each sample
  - Recommended plate layout (modify and record difference if necessary):
    - (“v1” stands for voltage setting 1, etc)
    - Correction: the medium and strong strains were swapped for both red and green fluorescent proteins. The plate below now explicitly provides tube labels. If you use a different layout (e.g. with swapped medium and strong strains), simply note the alternative layout.

| | 1 | 2 | 3 | 4 | 5 | 6 | 7 | 8 | 9 | 10 | 11 | 12 |
|---|---|---|---|---|---|---|---|---|---|---|---|---|
| A v1 | FF | PBS | FF | EV **T05** | FF | MG **T06** | FF | SG **T07** | FF | MR **T08** | FF | SR **T09** |
| B v1 | | | | | FF | bead | | | | | | |
| C v2 | FF | PBS | FF | EV **T05** | FF | MG **T06** | FF | SG **T07** | FF | MR **T08** | FF | SR **T09** |
| D v2 | | | | | FF | bead | | | | | | |
| E v3 | FF | PBS | FF | EV **T05** | FF | MG **T06** | FF | SG **T07** | FF | MR **T08** | FF | SR **T09** |
| F v3 | | | | | FF | bead | | | | | | |
| G | | | | | | | | | | | | |
| H | | | | | | | | | | | | |

  - Alternate recommended layout:
    - Correction: the medium and strong strains were swapped for both red and green fluorescent proteins. The plate below now explicitly provides tube labels. If you use a different layout (e.g. with swapped medium and strong strains), simply note the alternative layout.

| | 1 | 2 | 3 | 4 | 5 | 6 | 7 | 8 | 9 | 10 | 11 | 12 |
|---|---|---|---|---|---|---|---|---|---|---|---|---|
| A v1 | PBS | EV **T05** | MG **T06** | SG **T07** | MR **T08** | SR **T09** | | | | | | bead |
| B v2 | PBS | EV **T05** | MG **T06** | SG **T07** | MR **T08** | SR **T09** | | | | | | bead |
| C v3 | PBS | EV **T05** | MG **T06** | SG **T07** | MR **T08** | SR **T09** | | | | | | bead |
| D | | | | | | | | | | | | |
| E | | | | | | | | | | | | |
| F | FF | FF | FF | FF | FF | FF | | | | | | FF |
| G | FF | FF | FF | FF | FF | FF | | | | | | FF |

| | | | | | | | | | | | | |
|---|---|---|---|---|---|---|---|---|---|---|---|---|
| H | FF | FF | FF | FF | FF | FF | | | | | | FF |

This layout simplifies the pipetting and reduces the chance of errors in that process, but increases the complexity of the flow cytometer run specification and increases the chance of error in that process.

- Measure each sample at three voltages (low-low-low, med-med-med, high-high-high), in rising order of voltage: (total of 27 samples, 27 measurements, excluding fluid blanks before samples. Time should be <1 hour)
  - We want to standardize the number of events acquired by each instrument, and we've chosen to achieve this by standardizing the sample acquisition volume to 150uL. Please use whatever mechanisms are available for your machine to achieve this sample acquisition volume (e.g. by specifying the acquisition volume to your autosampler or manually stopping acquisition based on your instrument flow rate).
  - For each bacterial sample and for PBS blank, collect 150 uL and run full collected volume.
    - On the Attune that is 180 uL "total draw" and 150 uL "acquisition"
  - For each focusing fluid blank, collect 50 uL to check for and minimize carry-over.
  - Shoot for as close as you can to 60 uL/min flow rate
    - E.g., NIST: 100 uL/min; Rice, MIT: 60 uL/min
  - Record height, width, and area measurements for all channels.

#### Cytometer Tube Procedure:

- Prepare >= 20mL of 1X PBS filtered (0.2um). Confirm and record pH of the PBS that is used.
- Transfer 1mL of 1X PBS to cytometer tube.
- Prepare PBS-diluted cytometer tube samples.
  - Retrieve 1 Eppendorf tube of each strain from the -80C storage and 1 Eppendorf tube of calibration beads from the 4C storage.
  - Immediately place Eppendorf tubes containing strains in 42 degree water bath or heat block for exactly 60 seconds (do *not* place Eppendorf tube containing beads in water bath or heat block).
  - Add 990uL of 1X PBS to Eppendorf tubes containing 10uL of biological sample or 75uL of calibration beads. Vortex gently.
  - Transfer entire volume of PBS-diluted samples to individual cytometer tubes for each sample.
- Measure cytometer tube samples:
  - We want to standardize the number of events acquired by each instrument, and we've chosen to achieve this by standardizing the sample acquisition volume to 150uL. Please use whatever mechanisms are available for your machine to achieve this sample acquisition volume (e.g. by specifying the acquisition volume to your autosampler or manually stopping acquisition based on your instrument flow rate).
  - Shoot for as close as you can to 60 uL/min flow rate.

- Record height, width, and area measurements for all channels.
- Measure **1X PBS cytometer tube sample** until 150uL is collected. Repeat acquisition at all voltages determined in Procedure to Determine Voltages (the same cytometer tube sample is used for all 3 voltages, and the different voltages must be specified manually by the user).
  - At a flow rate of 60uL/min, acquire sample for 150 seconds (2 ½ minutes).
  - At a flow rate of 100uL/min, acquire sample for 90 seconds (1 ½ minutes).
- Measure each of the **PBS-diluted biological cytometer tube samples** until 150uL is collected. Repeat acquisition at all voltages determined in Procedure to Determine Voltages (the same cytometer tube sample is used for all 3 voltages, and the different voltages must be specified manually by the user).
  - At a flow rate of 60uL/min, acquire sample for 150 seconds (2 ½ minutes).
  - At a flow rate of 100uL/min, acquire sample for 90 seconds (1 ½ minutes).
- Measure **calibration bead cytometer tube sample** until 150uL is collected. Repeat acquisition at all voltages determined in Procedure to Determine Voltages (the same cytometer tube sample is used for all 3 voltages, and the different voltages must be specified manually by the user).
  - At a flow rate of 60uL/min, acquire sample for 150 seconds (2 ½ minutes).
  - At a flow rate of 100uL/min, acquire sample for 90 seconds (1 ½ minutes).

The above procedures for one replicate (Autosampler/Plate Procedure or Cytometer Tube Procedure) is to be repeated three times within three days. **It is OK to run multiple replicates back to back, just no slower than one per day.**

#### UD Sample Data Upload

- Data will be uploaded to the group through a Dropbox File Request link: [link removed]
- Please organize and name your files as follows:
  - Correction: Please label files by the **Tube ID** (not the Strain ID). Also, please label by Instrument ID (instead of Group).
  - [Instrument ID]/Replicate[N]/[Low|Med|High]/[sample abbreviation/Tube ID]_[well]_[#].fcs
    - If using tubes, drop well number
  - Examples:
    - MIT's 2nd replicate of "strong green" at high voltage in the recommended layout would be: N001/Replicate2/High/T07_E8_044.fcs
    - The focusing fluid blank after it would be: N001/Replicate2/High/T07_E9_045.fcs

        - Their beads for the 2nd replicate at low voltage would be: N001/Replicate2/Low/bead_B6_018.fcs
    - Complete pattern should have a collection of:
        - Autosampler: 7 Samples * 2 (plus blanks) * 3 voltages * 3 replicates = 126 files
            - [Instrument ID]/Replicate1/Low/FF_A1_001.fcs
            - [Instrument ID]/Replicate1/Low/PBS_A2_002.fcs
            - [Instrument ID]/Replicate1/Low/FF_A3_003.fcs
            - [Instrument ID]/Replicate1/Low/T05_A4_004.fcs
            - [Instrument ID]/Replicate1/Low/FF_A5_005.fcs
            - [Instrument ID]/Replicate1/Low/T06_A6_006.fcs
            - [Instrument ID]/Replicate1/Low/FF_A7_007.fcs
            - [Instrument ID]/Replicate1/Low/T07_A8_008.fcs
            - [Instrument ID]/Replicate1/Low/FF_A9_009.fcs
            - [Instrument ID]/Replicate1/Low/T08_A10_010.fcs
            - [Instrument ID]/Replicate1/Low/FF_A11_011.fcs
            - [Instrument ID]/Replicate1/Low/T09_A12_012.fcs
            - [Instrument ID]/Replicate1/Low/FF_B1_013.fcs
            - [Instrument ID]/Replicate1/Low/bead_B6_018.fcs
            - [Instrument ID]/Replicate1/Med/FF_C1_019.fcs
            - ...
            - [Instrument ID]/Replicate1/Med/bead_D6_036.fcs
            - [Instrument ID]/Replicate1/High/FF_E1_037.fcs
            - ...
            - [Instrument ID]/Replicate1/High/bead_F6_054.fcs
            - [Instrument ID]/Replicate2/Low/FF_A1_055.fcs
            - ...
            - [Instrument ID]/Replicate2/High/bead_F6_108.fcs
            - [Instrument ID]/Replicate3/Low/FF_A1_109.fcs
            - ...
            - [Instrument ID]/Replicate3/High/bead_F6_162.fcs
        - Tubes: 7 samples * 3 voltages * 3 replicates = 63 files
            - [Instrument ID]/Replicate1/Low/PBS_001.fcs
            - [Instrument ID]/Replicate1/Low/T05_002.fcs
            - [Instrument ID]/Replicate1/Low/T06_003.fcs
            - [Instrument ID]/Replicate1/Low/T07_004.fcs
            - [Instrument ID]/Replicate1/Low/T08_005.fcs
            - [Instrument ID]/Replicate1/Low/T09_006.fcs
            - [Instrument ID]/Replicate1/Low/bead_009.fcs
            - [Instrument ID]/Replicate1/Med/PBS_010.fcs
            - ...
            - [Instrument ID]/Replicate1/Med/bead_018.fcs
            - [Instrument ID]/Replicate1/High/PBS_019.fcs
            - ...

        - [Instrument ID]/Replicate1/High/bead_027.fcs
        - [Instrument ID]/Replicate2/Low/PBS_028.fcs
        - ...
        - [Instrument ID]/Replicate2/High/bead_054.fcs
        - [Instrument ID]/Replicate3/Low/PBS_055.fcs
        - ...
        - [Instrument ID]/Replicate3/High/bead_081.fcs
- Zip your directories into one big zip file, then upload that to the Dropbox request
    - Files uploaded to Dropbox request all arrive in the same directory, so making a zip file (or equivalent) is the only way to ensure your directory structure is preserved.
- Read-only link to data: [link removed]

#### A Analysis:

*Analysis only needs to be performed by the study organizers.*

##### Suggested Analysis Packages (listed alphabetically):

| Package Name | Documentation | Source Code |
|---|---|---|
| CytoFlow | [website] | [github] (Python) |
| FlowCal | [publication][website] | [github] (Python) |
| TASBE | [website] | [github] (MATLAB) |

##### Analysis Guidelines:

- Identify and align peaks, computing ERF/a.u. scaling factor per declared bead values
- For each biological sample:
    - Gate using a Gaussian mixture model on FSC and SSC to select the cellular component
    - Apply standard affine spectral compensation and background removal (e.g., per [Roederer '02]
    - Multiply all a.u. values by ERF/a.u. Scaling factor
    - Compute geometric mean of set of sample values above zero
- Measurands to compare:
    - Geo.std. of means across voltages within a sample set
    - Geo.std. of means across sample sets within a single lab
    - Geo.std. of means across laboratories
