## Supplementary material for "Meeting Measurement Precision Requirements for Effective Engineering of Genetic Regulatory Networks": S4: Flow Cytometry Data Processing

Flow cytometry data analysis was conducted using a unit conversion model from arbitrary units to MEFL/MEPTR constructed per the recommended best practices of TASBE Flow Analytics for each data set using the bead sample and lot information provided by each participating lab:

- Gating was automatically determined using a two-dimensional Gaussian fit on the forward-scatter area and side-scatter area channels for the first negative control. An example is shown in Figure 1.
- The same negative control was used to determine autofluorescence for background subtraction. Specifically, autofluorescence was estimated for each channel as a normal distribution (arithmetic mean and standard deviation of the gated negative control), and removed from channel values by subtracting the mean. An example is shown in Figure 2.
- As only one fluorescent protein was used in each strain, there was no need for spectral compensation or color translation.
- Calibration used standard SpheroTech Rainbow Calibration beads [1] for dye-based calibration to equivalent fluorescent molecules [2]. All beads were taken from the same samples of RCP-30-5A beads (lot AJ01) and conversion from arbitrary units to MEFL (for GFP strains) and to MEPE (for mCherry strains) was computed using the peak-to-intensity values provided by the manufacturer for those channels. Specifically, the peaks are fit against intensity values using a linear fit constrained to a slope of 1. Examples are provided in Figure 3 and Figure 4.

  Note that no instrument channel actually matched the laser/filter specifications for these channels, so these conversions were selected as the closest fit available. For GFP, the most common instrument channel (488 nm excitation, 530/30 bandpass filter) was close to the MEFL specification of 488 nm excitation, 530/40 bandpass filter. For mCherry, the most common instrument channel (561 nm excitation, 610/20 bandpass filter) does not match any laser specification for the beads, so we selected the MEPE specification of 488 nm excitation, 575/25 bandpass filter, which at least effectively matched two instruments having 488 nm excitation, 575/26 bandpass filter.

The color model thus produced for each data set was then applied to each sample in that data set to filter events and convert GFP and mCherry measurements from arbitrary units to MEFL and MEPE respectively, and geometric mean and standard deviation computed for the filtered collection of events.

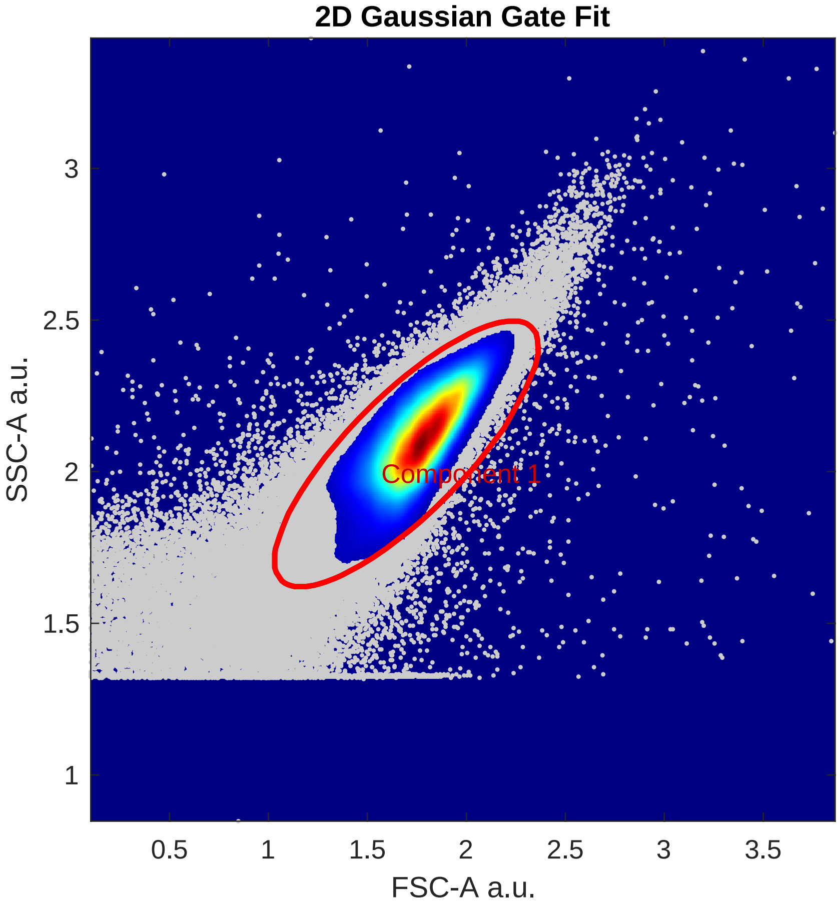


Figure 1: Prototypical example of two-dimensional Gaussian fit used for determination of gating from negative control. Points within the red circle are retained; points outside of the red circle are not.

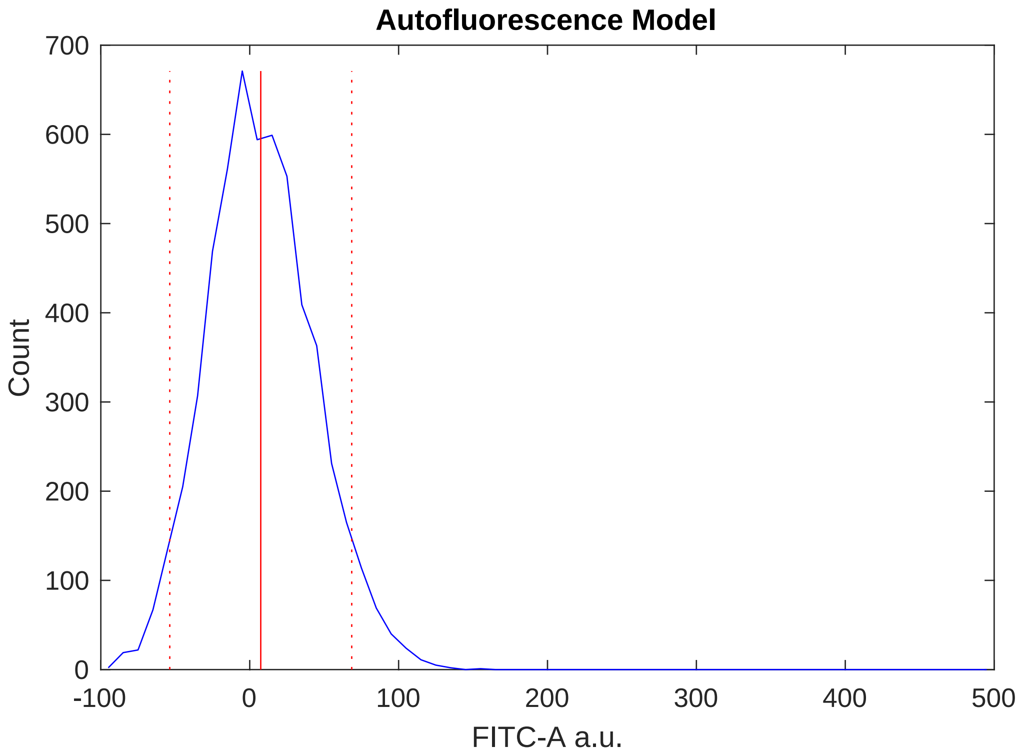


Figure 2: Prototypical example of mean and standard deviation computation of autofluorescence from negative control.

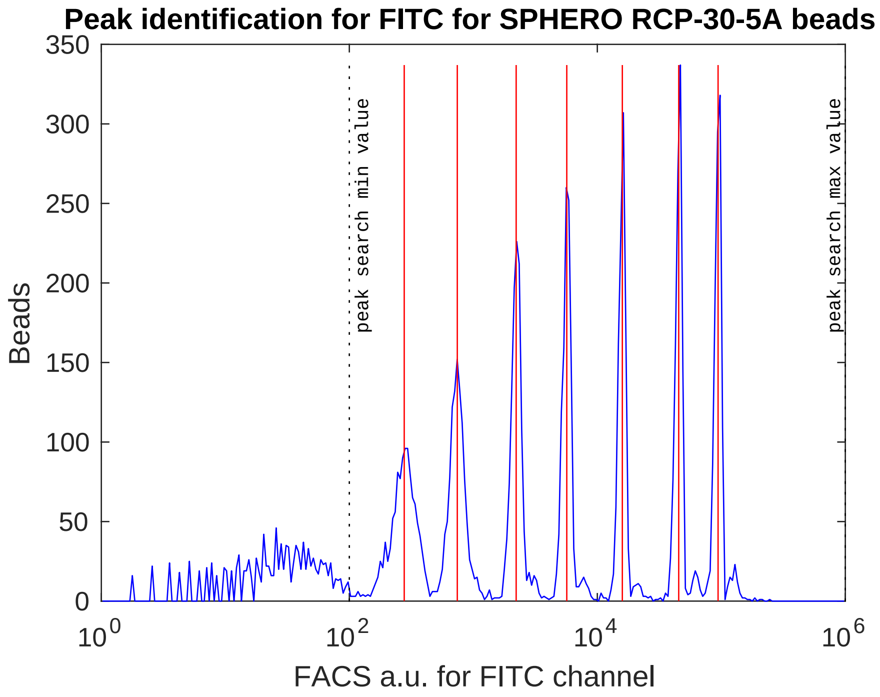


Figure 3: Prototypical example of calibration bead sample showing peak detection across a broad linear range.

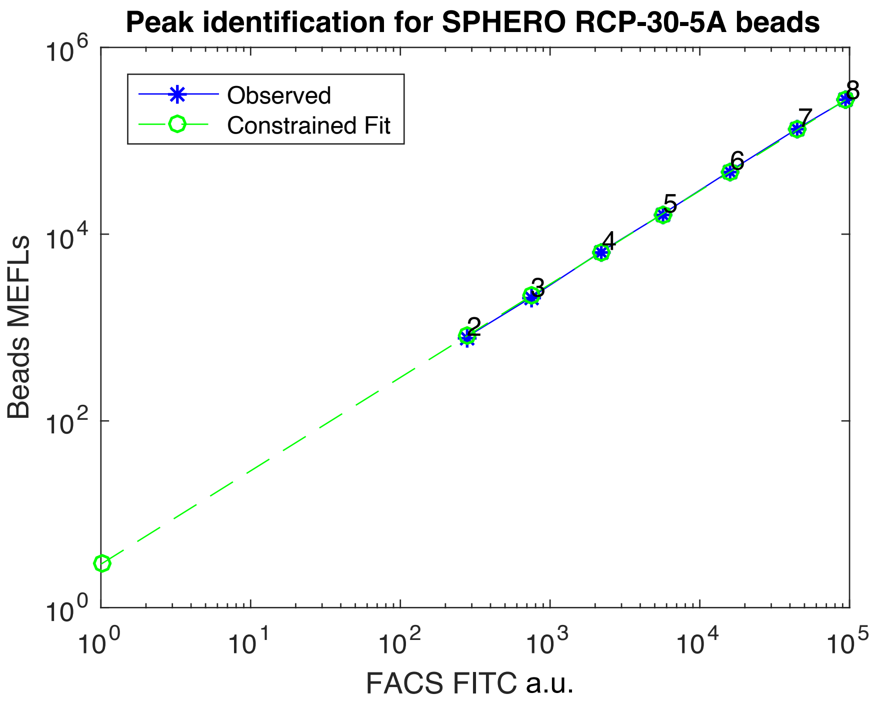


Figure 4: Prototypical example of unit conversion from arbitrary units to MEFL computed from calibration bead peaks.
